# UBiT2: a client-side web-application for gene expression data analysis

**DOI:** 10.1101/118992

**Authors:** Jean Fan, David Fan, Kamil Slowikowski, Nils Gehlenborg, Peter Kharchenko

## Abstract

We present a purely client-side web-application, UBiT2 (User-friendly BioInformatics Tools), that provides installation-free, offline alignment, analysis, and visualization of RNA-sequencing as well as qPCR data. Analysis modules were designed with single cell transcriptomic analysis in mind. Using just a browser, users can perform standard analyses such as quality control, filtering, hierarchical clustering, principal component analysis, differential expression analysis, gene set enrichment testing, and more, all with interactive visualizations and exportable publication-quality figures. We apply UBiT2 to recapitulate findings from single cell RNA-seq and Fluidigm Biomark™ multiplex RT-qPCR gene expression datasets. UBiT2 is available at http://pklab.med.harvard.edu/jean/ubit2/index.html with open-source code available at https://github.com/JEFworks/ubit2.

## INTRODUCTION

Recent rapid technological developments have enabled the generation of large amounts of biological data, which require computational analyses in order to draw meaningful conclusions and discover relevant biology. In particular, single cell transcriptome datasets, such as single cell RNA-sequencing data and Fluidigm Biomark™ multiplex RT-qPCR data, require analysis with methods beyond the capabilities of programs such as Excel. Although numerous R and Python packages, libraries, and scripts have already been developed to perform such analyses, alternative user-friendly, portable bioinformatics tools are still needed (Gehlenborg et al., 2010).

Web-based applications provide platform-independent, installation-free software solutions and have been historically popular for bioinformatics tools. Traditional web-based bioinformatics tools such Galaxy (Giardine et al., 2005; Goecks et al., 2010) depend on server back-ends to perform all computational analysis. Such server back-ends can be expensive to operate and maintain, and face challenges in speed and scalability due to server limitations. Less computationally intensive algorithms (Ca et al., 2014) and distributed computing between the server and client (Kim et al., 2014; Skinner et al., 2009) have been proposed as potential solutions to lessen server load. Still, issues of patient data privacy and confidentiality present additional concerns when data must be uploaded and stored on a server (Schatz et al., 2010). Recent advances in personal computing and HTML5-technology have enabled greater computational resources for client-side applications. Client-side applications address these server-limitation and privacy concerns since all computing is performed using the user's local computing resources such as their computer’s own memory and processor. Data is stored in memory locally and never uploaded onto an external server, enabling offline analysis like traditional downloadable stand-alone software but without the barrier of installations. Such technologies have already been applied to sequence alignment (Schmid-burgk and Hornung, 2015) and visualization (Yachdav et al., 2015), but have yet to be applied to gene expression data analysis through an integrated application. Here, we build on these advances to create UBiT2 (User-friendly BioInformatics Tools), lightweight, fully client-side set of web-based bioinformatics application for gene expression analysis.

## RESULTS

UBiT2 differs from existing systems in its lightweight, fully client-side implementation. As with other web-based bioinformatics applications, UBiT2 uses the same standard algorithms and statistics that users can typically accomplish in R (Figure 1, Supplementary code). However, its fully client-side implementation in JavaScript means your data is never accessed or stored on an external server, and all computation and analyses are executed by the user’s Internet browser, using the user’s own computer’s memory and CPU. Current analysis modules have been designed for single cell gene expression analysis starting from either from an expression matrix (Figure 2), or from FASTQ files (Figure 3). Users select their FASTQ files or copy and paste in their gene expression matrix to perform all alignments and downstream analysis such as principal component analysis (PCA), hierarchical clustering, differential expression analysis, and more. UBiT2’s flexibility of data input, workflow, and seamless integration of read-mapping and quantification with the standard downstream analyses enables quick and lightweight single cell gene expression analysis. Interactive visualizations enable rapid exploration of the impact of data filtering and method parameters. Both presentation-ready and publication-ready figures in PNG and SVG formats can be readily exported.

**Figure 1.**
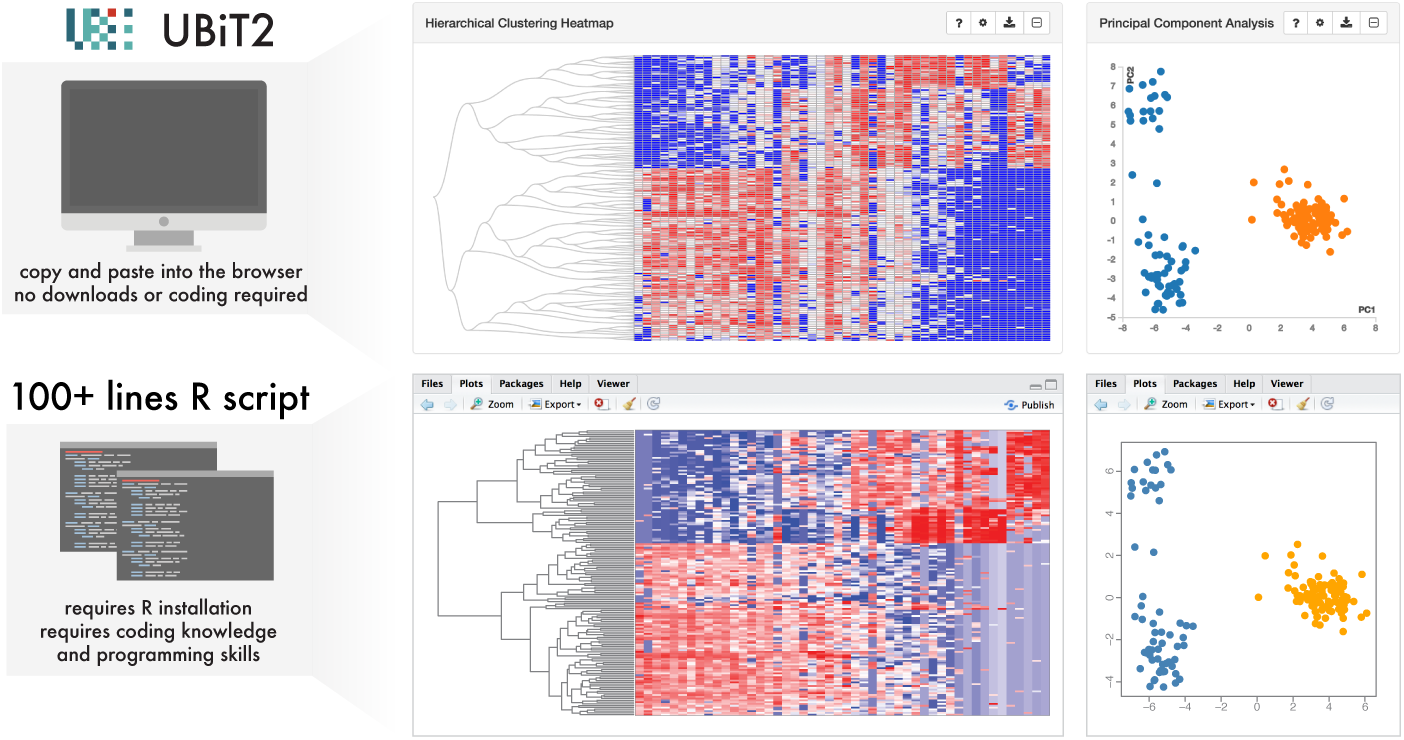
Sample analysis using UBiT2 compared to 100+ lines of R code. Results for hierarchical clustering, principal component analysis, and K-means clustering are virtually identical, as expected. Thus the same results and figures can be achieved through UBiT2 with coding knowledge and programming skills.

**Figure 2.**
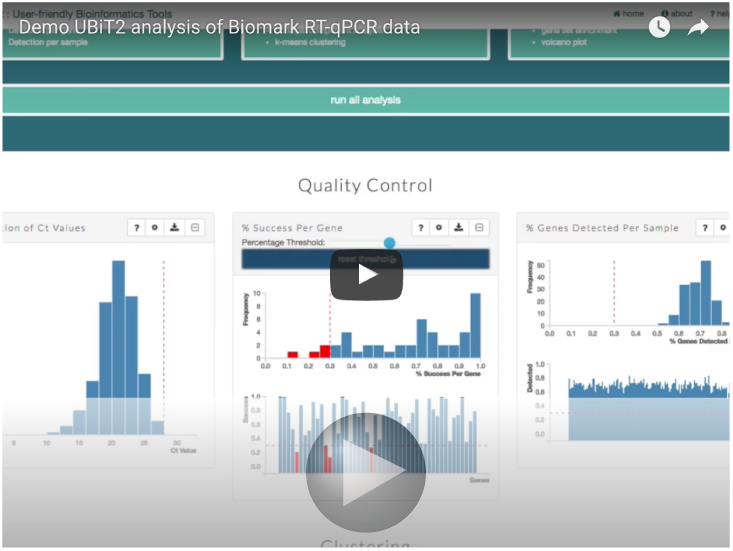
Video tutorial of analysis with UBiT2 starting from a gene expression matrix. As demonstrated in the video, users can select to start from a gene expression matrix. A gene expression matrix can be copied directly from Excel into the UBiT2 interface. The user can interactive toggle options using the gear icon and rerun analyses as needed.

**Figure 3.**
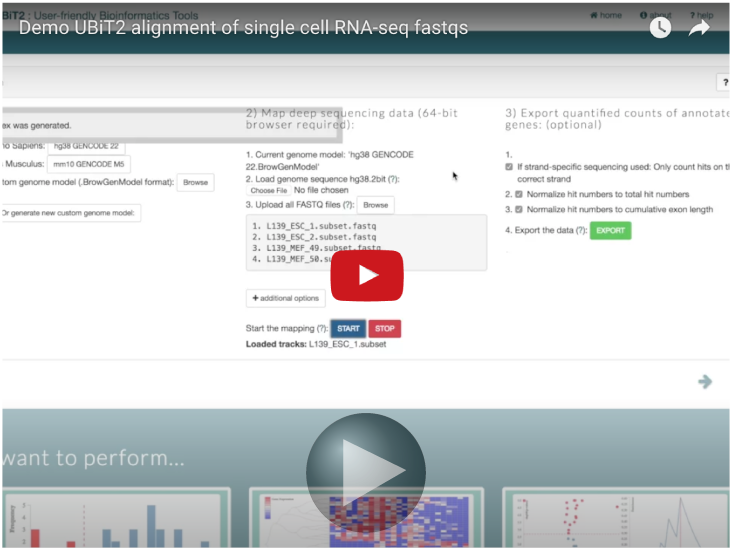
Video tutorial of alignment and expression quantification with UBiT2 starting from FASTQ files. As demonstrated in the video, users can select to start from FASTQ files. Users select their reference genome or upload their own. Users can upload up to 96 FASTQ files by simultaneously selecting multiple files. Results are then automatically ported into the expression matrix tab and can be downloaded.

### UBiT2 accurately identifies and characterizes transcriptional subpopulations using qPCR data

We apply UBiT2 to identify and characterize transcriptional subpopulations in the 64-cell stage blastocyst using Fluidigm Biomark™ multiplex RT-qPCR data from Guo et al. (2010). After downloading the original Excel spreadsheet for 159 cells collected from the 64-cell stage blastocyst, we copy and paste the data into the gene expression matrix table in UBiT2. We then applied an LoD method transformation and set a Ct value of 28 as the failure threshold, as recommended by Livak et al. (2013) for Biomark™ data. The resulting principal component analysis quickly identifies three subpopulations (Figure 4A), recapitulating published findings (Guo et al., 2010). Results from the differential expression analysis between group 1 (blue) and group 2 (orange) is visualized as a volcano plot (Figure 4B), highlighting *Id2* and *Pdgfra* as two significantly differentially expressed genes with large fold changes. Plotting each gene as a beeswarm plot allows us to further assess the distribution of gene expression within each identified subpopulations (Figure 4C) to confirm that *Id2* is indeed more highly expressed in group 2 while *Pdgfra* is more highly expressed in group 2. Further interactive exploration of known markers such as *Cdx2*, *Nanog*, and *Gata4* allows us to then define our identified subpopulations as trophectoderm, epiblast, and primitive endoderm (Guo et al., 2010). In contrast, house-keeping genes such as *Gapdh* do not show substantial expression differences between identified subpopulations as expected (Figure 4C).

**Figure 4.**
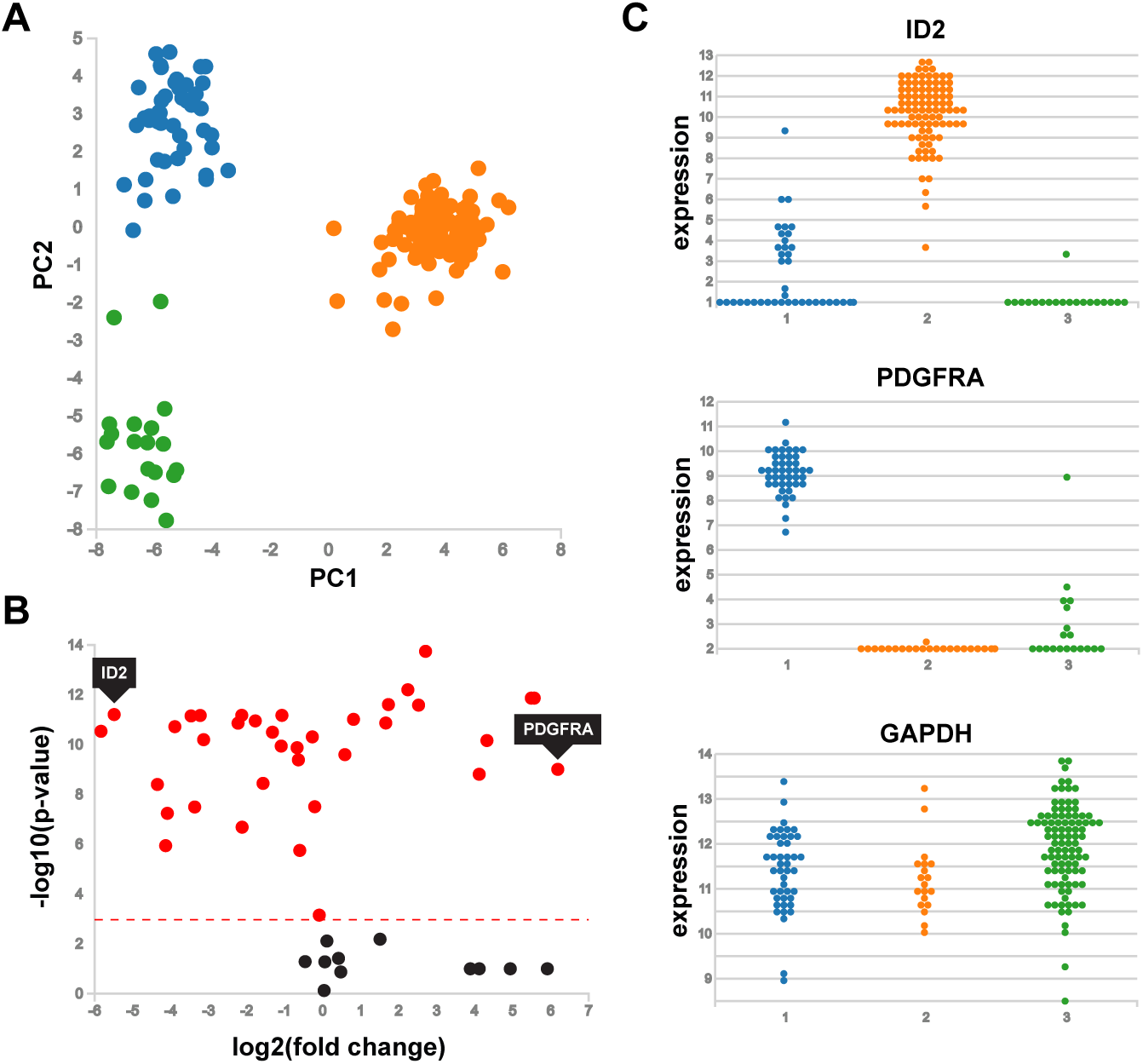
Analysis of Fluidigm Biomark™ multiplex RT-qPCR data for 159 cells collected from the 64-cell stage blastocyst from Guo et al. (2010) using UBiT2. A) Principal component analysis identifies 3 transcriptionally distinct cell subpopulations. Each point is a single cell. Cells are colored by labels from K-means clustering with K=3. B) Volcano plot of differential expression analysis results comparing group 1 vs. group 2. Each point is a gene. The horizontal line denotes the significance threshold of p-value = 0.05. C) Beeswarm plots comparing gene expression in each identified cell subpopulation. Each point is a single cell. Cells are grouped and colored by labels from K-means clustering.

### UBiT2 accurately identifies and characterizes transcriptional subpopulations using single cell RNA-seq data

We also apply UBiT2 to identify and characterize transcriptional subpopulations in 226 fetal cortical single cells from Camp et al. (2015). Visualizing the distribution of zeros (Figure 5A), we can see a clear zero-inflation in this dataset. Considering all zeros as ‘failures’, we can then visualize the distribution of detection per gene and genes detected per sample to assess data quality (Figure 5B). Principal components analysis with K-means clustering with k=2 along with hierarchical clustering identifies two main transcriptional subpopulations. Gene set enrichment analysis on genes ranked by fold-change identifies Neuronal Differentiation as a signficantly enriched gene set (Figure 5C), suggesting that subpopulations may correspondend to neuroprogenitors versus mature neurons, consistent with published findings (Camp et al., 2015). Indeed, visualization of known the neuronal growth and maturation marker *STMN2* further confirms subpopulation identities (Figure 5D).

**Figure 5.**
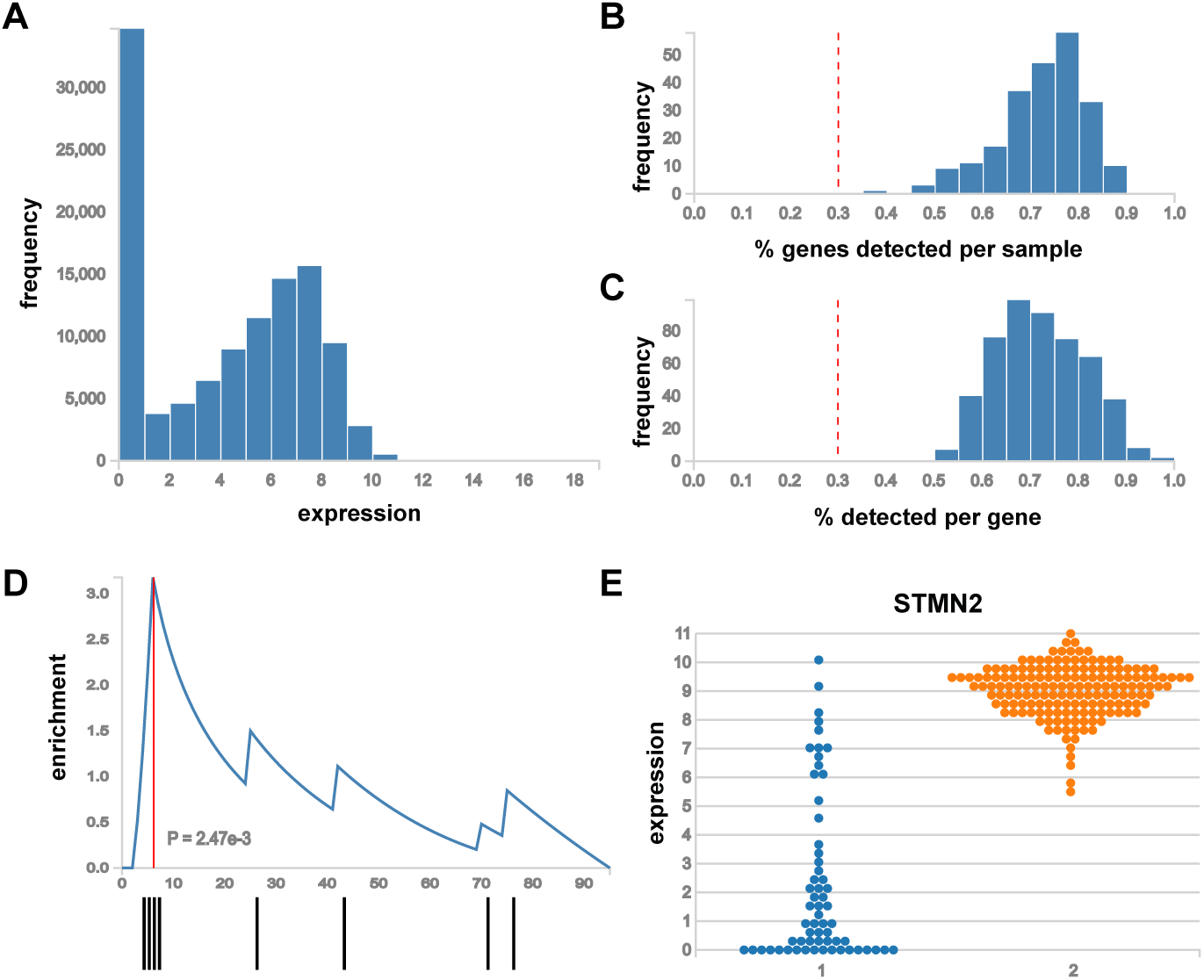
Analysis of single cell RNA-seq data for 226 fetal cortical single cells from Camp et al. (2015) using UBiT2. A) Histrogram of expression. B) Histogram of the percentage genes detected per sample with detection meaning non-zero expression. C) Histogram of the percentage detected per gene. D) Gene set enrichment plot for GO:0030182 (neuron differentiation). Genes sorted by p-value. Each line (bottom) means a gene at that rank is within the gene set. Red line denotes the point of highest enrichment. P-value obtained by non-parametric minimal hypergeometric test XL-mHG (Eden et al., 2007; Wagner, 2016).

## DISCUSSION

As the size of datasets in the biological domain increases, the need emerges for scalable bioinformatics analysis services that are easy for all to use. Here, we present UBiT2 to provide such common bioinformatics analysis for gene expression data.

We envision future developments in bioinformatics web-applications such as UBiT2 will enable direct and seemless integration with public data. For example, The Gene Expression Omnibus (GEO) provides over 6,000 curated gene expression datasets, including gene expression values and sample annotations (Edgar et al., 2002; Barrett et al., 2013). In addition, bioinformatics experts need better tools to create interactive reports to share with collaborators. Future work on UBiT2 will include an export feature to generate a standalone HTML file that includes sample annotations, gene expression data, and all of the Javascript code for analysis and visualization. Since many datasets are just a few megabytes in size, this standalone HTML file will be easily shareable by email. All collaborators, regardless of programming experience, will be empowered to explore their datasets in this interactive report.

Although UBiT2 is neither designed to perform full in-depth analyses, nor is it designed to replace server-side high-performance computing, it serves to provide users with a preliminary look into their data that is convenient and private for the user and cost-effective and highly scalable for the developer.

## METHODS

All analysis and visualization components are built on web standards, including JavaScript, CSS, and HTML. Visualizations are built on D3.js Bostock et al. (2011) and additional APIs are used to interface with databases such as mygene.info (Wu et al., 2013).

### Data

The gene expression matrix for Guo et al. (2010) was loaded from the supplementary material of the original paper (mmc4.xls). Data corresponding to the 64 cell sample (cell names prefixed with 64C) was selected for analysis.

The FPKM matrix for Camp et al. (2015) was downloaded from GEO under accession GSE75140. For demonstrative purposes, data was parsed using R to restrict to only the 226 fetal cortical cells and filtered for 500 highly expressed and variable genes (See Supplementary code).

To assist users with using UBiT2, a small dataset from Lawson et al. (2015) is provided by default.

### Data transformations

Data is expected to be in the format of cells or samples as rows and genes as columns. Data can be transposed by checking the transpose box. Data can be transformed according to the following options: 1. none, 2. LoD, 3. log10, or 4. asinh.

### Quality control and filtering

By default, expression entries denoted as 999, the default failure value for the heatmap export from the Fluidigm Biomark™ platform, will be considered detection failures. Percentage detection per gene and percentage genes detected per sample can thus be calculated as the percentage of non-failures per column and per row respectively. Genes and samples with a greater percentage detection than a user-specific threshold, by default 30%, will be used in downstream analysis. Users can adjust either threshold using the sliding threshold bar.

### Alignment and Mapping

UBiT2 incorporates BrowserGenome.org’s ability to perform client-side alignment and quantification (Schmid-burgk and Hornung, 2015). Briefly, BrowserGenome.org is a web-based read mapping and transcript quantification web tool implemented in Javascript. It utilizes a non-overlapping q-gram indexing algorithm with sorted data structures and random sampling to enable quantification of gene expression in the web browser. Because BrowserGenome’s read-mapping strategy is optimized to reduce the computational load for quantification, it is able to match the performance of established software such as STAR in both speed and accuracy. In the same vein as BrowserGenome, users of UBiT2 may choose a default reference genome or create a new one with a .2bit file and GENCODE .gtf gene model. We extend BrowserGenome’s capabilities by allowing the user to upload up to 96 FASTQ files at once instead of one file at a time, and having UBiT2 read-map and quantify the files in a continuous pipeline. Index generation progress is shown dynamically as a percentage for the current file, and the overall progress of the read-mapping and quantification process is displayed visually with a progress bar. Upon completion, the user may then export the quantified expression or continue on UBiT2 with downstream bioinformatics analyses.

### Clustering

UBiT2 uses a standard bottom-up agglomerative hierarchical clustering with a modified version of clusterfck (https://harthur.github.io/clusterfck to cluster samples. Users can select to specify the distance metric (Euclidean, Manhattan, max, or correlation-based) and the linkage criterion (average, single, or complete). Alternatively, users can cluster data by applying dimensionality reduction using principal component analysis prior to k-means clustering. Users specify k.

### Differential expression testing

Expression between groups identified by clustering can be compared using a Wilcoxon rank sum test. Users specify the two groups to be compared.

### Gene set enrichment analysis

UBiT2 uses a novel, JavaScript implementation of the non-parametric minimal hypergeometric test XL-mHG (Eden et al., 2007; Wagner, 2016) to perform gene set enrichment testing. Users have the option of ranking genes by p-value or fold-change. User-specified rankings also determine the row orderings of the heatmap visualization. An optimized and streamlined set of gene set annotations is used to assist with gene to gene set mapping (Carlson, 2015). An API is used to interface with as mygene.info(Wu et al., 2013) to provide gene information. Current gene set enrichment analysis and gene information lookup supports gene names for Homo sapiens as symbols consistent with HGNC nomenclature (Gray et al., 2015).

## ACKNOWLEDGEMENTS

This research was supported by the Ruth L. Kirschstein National Research Service Award from the National Cancer Institute for JF (F31CA20623), and the National Institute of Arthritis and Musculoskeletal and Skin Diseases for KS (F31AR070582), the NIH Department Of Health And Human Services (R00HG007583) for NG, and the NIH Centers for Excellence in Genomic Science (P50MH106933) and NSF CAREER (NSF-14-532) awards for PVK.

